# Mobile genetic elements drive a fusion-deletion life cycle that shapes plasmid evolution and antimicrobial resistance

**DOI:** 10.64898/2026.01.09.696371

**Authors:** Thomas Ipoutcha, Yiqing Wang, Eduardo P C Rocha, José R Penadés

## Abstract

Plasmids are key drivers of bacterial adaptation, yet the mechanisms that generate their diversity remain poorly understood. Here, we show that mobile genetic elements (MGEs) orchestrate a fusion-deletion life cycle that repeatedly remodels plasmids during evolution in *Staphylococcus aureus*. Large-scale genomic analyses reveal that multireplicon plasmids are widespread and strongly enriched in transposases. Using experimental assays, we demonstrate that rare MGE-mediated fusion events, via homologous recombination or transposition, combine distinct plasmids into single multireplicon elements, expanding gene content and transfer potential. Antibiotic pressure selectively enriches these fused plasmids, rescuing bacterial populations under stress, whereas opposing selective forces, including phage predation, favour deletion derivatives that preserve essential functions and phage transmissibility. This cyclical process generates dynamic plasmid repertoires with conserved backbones and diverse accessory modules. We propose that MGE-driven fusion-deletion cycles represent a general principle of plasmid evolution, explaining the rapid emergence and persistence of multidrug-resistant plasmids across bacterial pathogens.

## Introduction

Plasmids are central drivers of bacterial evolution, mediating the horizontal transfer of genes that confer antibiotic resistance, virulence, and metabolic versatility. ^1,2^ Their mobility enables rapid adaptation across ecological and clinical contexts and has been a major force in the emergence and global spread of multidrug-resistant pathogens, particularly among the ESKAPE bacteria. ^3^ As antimicrobial resistance (AMR) continues to escalate worldwide, ^4,5^ identifying the evolutionary principles that govern plasmid diversity has become a critical priority.

Across bacteria, plasmids adopt distinct mobility strategies, ranging from large self-transmissible elements (conjugative) to smaller mobilizable plasmids that rely on other mobile elements for dissemination. ^6^ Conjugative plasmids encode the full machinery required for self-transfer, whereas mobilizable plasmids carry only partial transfer functions, such as an origin of transfer and sometimes a relaxase, and depend on a conjugative plasmid to move. ^7^ Non-mobilizable (also referred to as non-transmissible) plasmids depend on phage-mediated transduction for their transfer, while some plasmids (around 10% in *Staphylococcus aureus*) lack any known transfer mechanism. Conjugative plasmids are generally large (>50 kb, > 25kb in firmicutes), ^8^ reflecting the need to encode the genes for the mating pair formation system. In contrast, across all analysed species, mobilizable and non-transmissible plasmids display a bimodal size distribution, ^1^ with peaks corresponding approximately to the size of temperate phage genomes (∼40–45 kb) and phage satellites (∼10 kb) in *Staphylococcus*. ^9,10^ This bimodality explains how smaller plasmids lacking conjugation-related genetic elements can nevertheless be mobilised via phage- or satellites-mediated generalised transduction. ^11^

These mechanisms of transfer impose strong physical and ecological constraints on plasmid size, creating a fundamental trade-off between mobility and gene content. ^1^ Smaller plasmids can be efficiently transferred by phages but encode limited accessory functions, whereas larger plasmids carry extensive adaptive cargo but rely almost exclusively on conjugation. Despite these constraints, natural plasmid populations display remarkable diversity in size, structure, and gene content, raising a central unresolved question: how is plasmid diversity generated and maintained in the face of strong selective and physical limitations? Although plasmids can gain, lose, and rearrange genetic material, the evolutionary logic governing these processes remains poorly understood. ^12^

Multireplicon plasmids have been described for decades and result from plasmid fusion events. Such plasmids have been observed in several pathogenic genera, including *Klebsiella*, ^13–16^ *Acinetobacter*, ^17^ *Salmonella*, ^18^ *Citrobacter*, ^19^ and *Staphylococcus*. ^20^ Historically, multireplicon plasmids were largely regarded as rare or exceptional outcomes of plasmid evolution, documented mainly as isolated case studies rather than as a pervasive evolutionary strategy.

Recent comprehensive bioinformatic analyses have revealed that multireplicon plasmids are widespread and play a major role in the emergence of clinically important plasmids. These studies have shown that plasmids have progressively accumulated antimicrobial resistance (AMR) genes, ^21–23^ and that multireplicon plasmids have become especially prevalent since the introduction and widespread use of antibiotics. ^22^ However, despite growing recognition of their importance, the prevalence, evolutionary significance, and mechanistic basis of plasmid fusion remain poorly resolved in many bacterial species, including *Staphylococcus aureus*, and it is unclear how rare fusion events become evolutionarily significant, how they are integrated over time, or whether multireplicon plasmids represent stable endpoints or transient intermediates in plasmid evolution.

Here, we propose that plasmids follow a defined life cycle that operates at the population level and is driven by mobile genetic elements. In this life cycle, rare fusion events transiently expand plasmid size, gene content, and mobility, whereas opposing selective forces, including bacteriophages, promote deletion and resolution, eliminating excess DNA while preserving essential functions and transmissibility. Unlike the life cycles of bacteriophages or other mobile elements, which largely preserve element identity across generations, this plasmid life cycle is inherently diversifying. Fusion generates genetic novelty, and subsequent deletion or resolution produces multiple distinct descendants rather than a single reverting form. This recurrent remodelling, rather than strict identity preservation, enables plasmid populations to explore a broad genetic space while maintaining conserved functional architectures. Our results therefore provide a mechanistic explanation for how plasmids achieve extensive genetic diversity despite strong constraints on size and mobility, reconciling rapid adaptation with long-term evolutionary stability.

## Results

### Plasmid diversity and fusion signatures in *S. aureus*

To determine how plasmid diversity is generated and maintained despite strong constraints on size and mobility, we analysed plasmids from *Staphylococcus aureus*, a species in which plasmid-mediated gene transfer is central to adaptation, virulence, and antimicrobial resistance. ^2^ The coexistence of conjugative, mobilizable, and non-transmissible plasmids within the same populations provides a natural system to assess how different mobility strategies, genetic architectures, and fusion signatures are distributed at scale.

We compiled a dataset of 2,564 complete *Staphylococcus* plasmids from the PLSDB database, including 1,649 from *S. aureus*. Plasmids were classified according to predicted mobility and replicon content (Fig. 1). Among plasmids carrying at least one identifiable Rep gene (2,308/2,564), 7% were conjugative (159), 22% mobilizable (516), 43% encoded an origin of transfer (997), and 31% were non-transmissible (636) (Fig. 1A).

**Figure 1:**
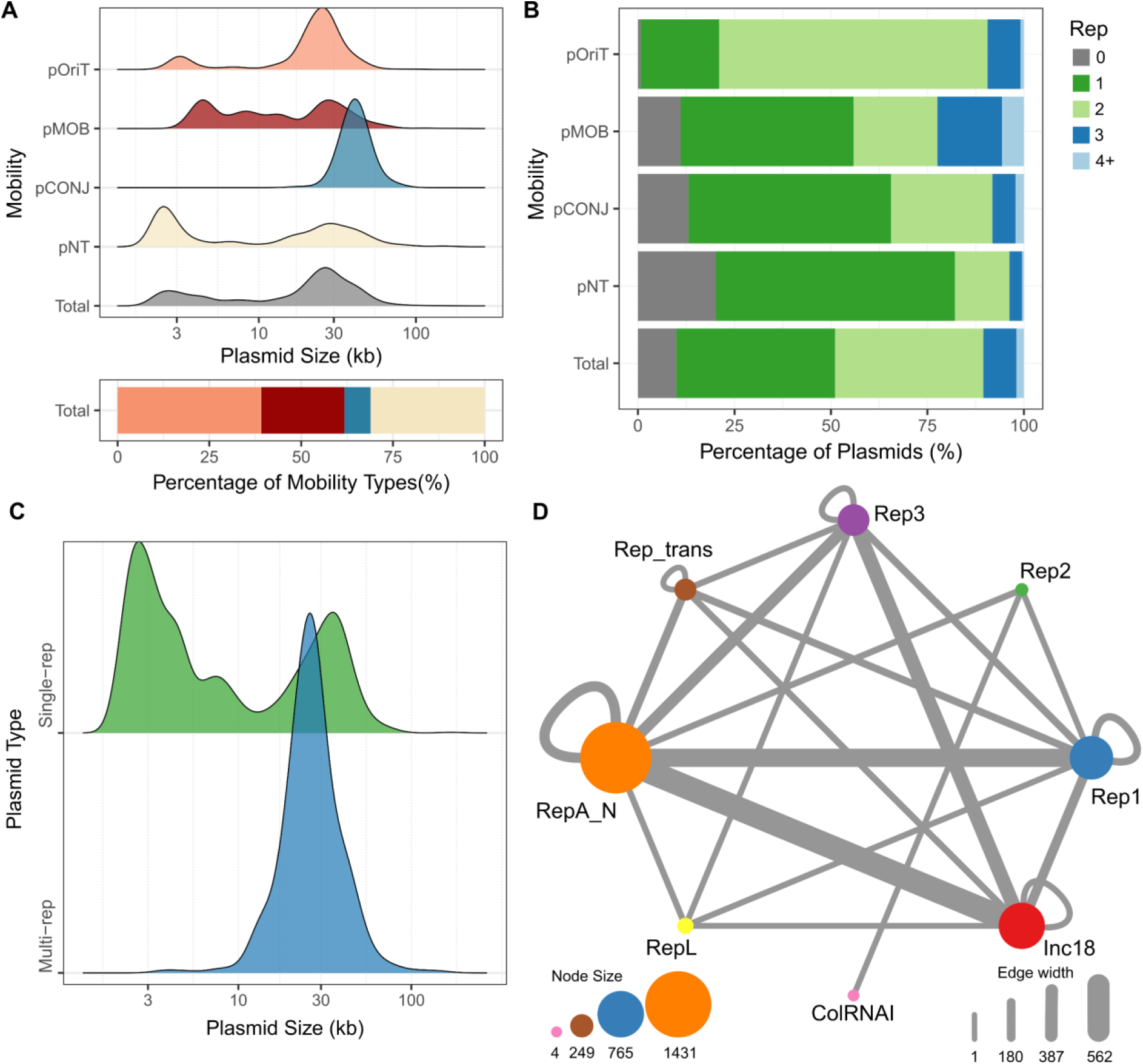
Analysis of mobility and replicon types of 2,564 complete *Staphylococcus* plasmids. pConj: Conjugative, pMob: mobilizable, pOriT: carrying oriT, pNT: Non-transmissible. **A-** Top: Density plot of plasmid size (log10 transformed) by mobility type (pOriT, pMOB, pCONJ and pNT). Bottom: Proportion of plasmids assigned to four mobility types (same color palette in top). **B-**Proportion of plasmids encoding 1-4 replicon(s) by mobility types, plasmids with more than four replicons are merged into “4+” group, plasmids encoding replicon not identifiable are in group “0”. **C-** Size comparison between single- and multi-replicon plasmids. **D-** Replicon co-occurrence network for multi-replicon plasmids. Nodes represent plasmid replicon families; node size is proportional to the number of replicons. Edges and self-loops indicate co-occurrence of two replicons from different and the same families on the same plasmid; edge thickness reflects the number of co-occurrence instances. Rolling-circle replicon: Rep1, Rep2, RepL, Rep_trans; theta replication: RepA_N, Inc18, Rep3, ColRNAI.

Unexpectedly, nearly half of all plasmids (49%, 1,255/2,564) carried more than one Rep gene, indicative of plasmid fusion events and the formation of multireplicon plasmids (Fig. 1B). Fusion signatures were particularly enriched among plasmids that rely on conjugation for transfer, whereas they were substantially less frequent among non-transmissible plasmids (Fig. 1B). Consistent with plasmid co-integration, multireplicon plasmids were significantly larger than single-replicon plasmids (Fig. 1C), indicating that fusion contributes directly to plasmid expansion.

Analysis of replicon families revealed that the presence of multiple replicons within a single plasmid, considered a hallmark of plasmid fusion, is widespread and encompasses nearly all possible combinations of replicons (Fig. 1D; Fig. S1A). Small plasmid families, such as those carrying the RepL and Rep_trans replicon, were typically restricted to sizes of 2–8 kb unless incorporated into larger fused plasmids (Table S1, Fig S1C), but RepL replicons were also present in plasmids with other replicons. Plasmids encoding RepA_N, Rep3 and Inc18 families showed significantly higher frequencies of fusion (Fisher’s exact test, p-value < 0.001), suggesting that certain replicon types play a central role in plasmid co-integration and diversification. Interestingly, plasmids with such replicons tend to be larger and mobilizable by conjugation (Figure S1B). Together, these results show that plasmid fusion is frequent in *S. aureus*, providing a potential mechanism for generating genetic diversity.

### Association of mobile genetic elements with plasmid fusion

Plasmid fusion is favoured by the presence of identical or highly similar sequences in distinct plasmids. Because insertion sequences (IS) are involved in plasmid evolution and are overrepresented in plasmids of clinical importance, ^24,25^ transposases are strong candidates for mediating such co-integration events. Indeed, IS elements, particularly those of the IS*6* family (e.g., IS*257*), were shown to generate transient plasmid co-integrates in *Escherichia coli* ^26–30^ and *Staphylococcus aureus*. ^20,31,32^ To determine whether IS-mediated fusion is a widespread feature of plasmid populations, we analysed the *Staphylococcus* plasmid dataset for the presence of insertion sequences.

BLASTn analysis revealed that 40% of *Staphylococcus* plasmids carry IS*257*, with a significant enrichment among multireplicon plasmids (Fisher’s exact test, p-value<0.001. 58% in plasmids with two replicons and up to 98% in plasmids with four replicons, compared with 22% in single-replicon plasmids) (Fig. S2B). Conjugative and *oriT*-encoding plasmids, both of which display elevated fusion frequencies, were also enriched for IS*257* (52% and 50%, respectively) (Fig. S2C). In contrast, RepL plasmids showed low frequencies of both IS*257* carriage (12%) and fusion signatures (10%) (Fig. S1A, S2A). Conversely, replicon families with high fusion prevalence, such as RepA_N, were strongly enriched for IS*257* (67%).

The Mu-family transposase encoded in IS*Bli29* (hereafter IS*Bli29* ORF4 transposase referred as IS*Bli29*), was one of the most abundant IS in *Staphylococcus* plasmids and showed a similar association with plasmid fusion. IS*Bli29* was rare in single-replicon plasmids (3%) but enriched in multireplicon plasmids (18–26%) (Fig. S2B). Notably, IS*Bli29* was particularly prevalent in Rep3 plasmids (39%), a family in which IS*257* elements are less common (27%) (Fig. S2A) yet fusion frequencies are high (77%) (Fig. S1A). Both IS*257* and IS*Bli29* were associated with increased plasmid size (Fig. S2D), consistent with their role in promoting plasmid co-integration. ^25,33^

We then assessed the impact of plasmid mobility, size and presence of ISs on the probability that the plasmid is multi-replicon using a generalized linear model and found that all variables are contributing significantly and in a positive manner for the later, i.e. multi-replicon plasmids tend to have more ISs, larger size and are more likely to be mobile or mobilizable by conjugation (Table 1). Together, these results identify transposable elements as central drivers of plasmid fusion, providing a mechanistic link between the widespread occurrence of multireplicon plasmids and the enrichment of insertion sequences in plasmid genomes.

**Table 1:**
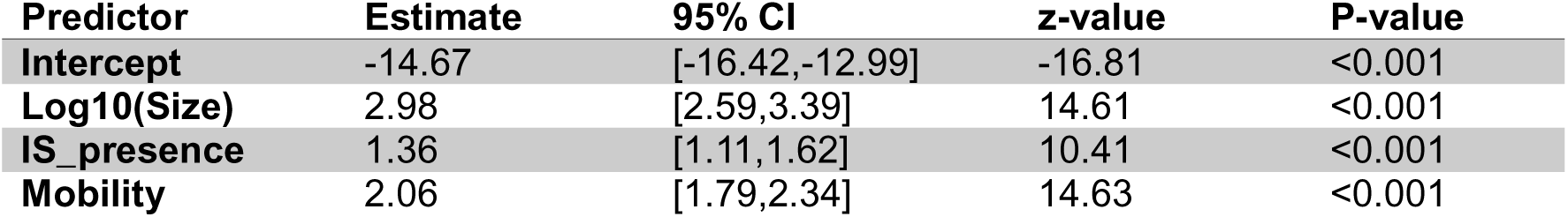
Binomial generalized linear model for multi- / single-replicon plasmid. Plasmid encoding multiple- / single-replicon was analyzed using a binomial glm with a logit link. Plasmid size shows siginificant effect, IS_presence (presence of either IS*6* or IS*Bli29*) also had an independent significant effect. Compared to plasmid lacking mobility (pNT), pOriT and pMOB show significant effect, pCONJ shows moderate effect.

### Plasmid families with conserved backbones show evidence of modular evolution

To investigate how plasmids diversify while maintaining conserved core functions, we asked whether plasmids sharing the same replicon type (the same replication machinery), also vary in size and gene content. If so, this would indicate that plasmids undergo repeated gain and loss of accessory genes over time. We focused on plasmids carrying the RepA_N replicon, which display a high prevalence of multireplicon architectures (Fig. S1A). Within this group, we analysed two representative subfamilies, Rep15 and Rep20 (Fig. 2A), both of which frequently carry IS*257* (Fig. 2B) and exhibit substantial size variability (Fig. 2C).

**Figure 2:**
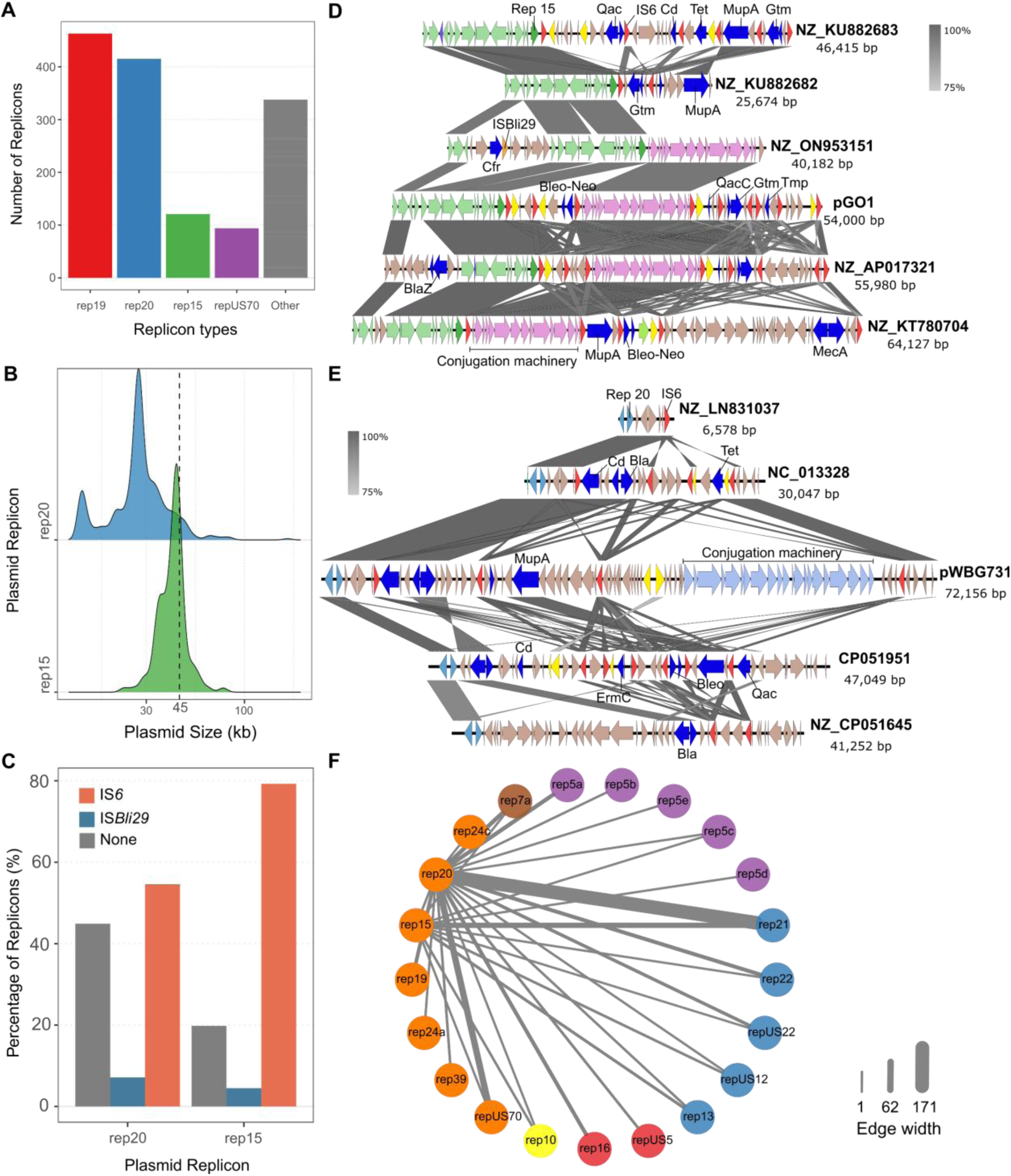
Rep15 and Rep20 plasmids. **A-** Replicon types in RepA_N family. (Rare replicon types are grouped into ‘Other’) **B-** Size comparison between plasmids encoding rep15 and rep20 replicon. **C-**Proportion of IS*6* and IS*Bli29* in plasmids encoding rep15 and rep20. **D**-Plasmid alignment of selected examples for Rep 15 plasmid family. **E-** Plasmid alignment of selected examples for Rep 20 plasmid family. Replication genes are colored in yellow, insertion sequence genes 6 are colored in red, AMR genes are colored in blue. Gtm: gentamycin resistance, Cd: Cadmium resistance, Tmp: Trimethoprim resistance, Erm: Erythromycin resistance. Bleo: Bleomycin resistance, MupA: Mupirocin resistance, Bla: beta-lactam resistance and Qac: quaternary ammonium compounds resistance. **F**-Replicon co-occurrence network for plasmids encoding rep15 and rep20. Nodes represent plasmid replicon types, colored by replicon families (the same color palette in Fig. 1, Theta replicon: RepA_N in orange, Inc18 in red, Rep3 in purple; Rolling-circle replicon: Rep1 in blue, Rep2 in green, RepL in yellow, Rep_trans in brown). Edges indicate co-occurrence of two replicon type on the same plasmid; edge thickness reflects the number of co-occurrence instances.

Comparative sequence analyses revealed highly conserved backbone regions within each subfamily, contrasted by important variation in accessory content and particular in AMR genes (Fig. 2D, E). Many accessory modules corresponded to complete or partial plasmid sequences flanked by IS*257* elements, often containing additional ISs, suggesting a role for transposases in both the acquisition and rearrangement of these modules. In the Rep15 subfamily, 90% of plasmids retained a conjugation module, whereas a minority lacked this region, consistent with deletion events (Table S1; Fig. 2D). Conversely, Rep20 plasmids were predominantly *oriT*-encoding or mobilizable, but a small subset carried conjugation machinery, consistent with sporadic acquisition events (Table S1; Fig. 2E).

Despite frequent fusion and rearrangement, plasmid sizes in both Rep15 and Rep20 remained restricted, with very few exceeding 45 kb (Fig. 2C). This pattern extended to the full *Staphylococcus* plasmid dataset, where only 6% of plasmids (162/2,564) were larger than 45 kb (Table S1).

Together, these observations indicate that plasmids with conserved replication backbones undergo continuous, modular remodelling through the gain and loss of accessory regions. This process can generate important diversity in antimicrobial resistance gene combinations while preserving core plasmid functions. The mechanisms underlying this modular evolution are explored in the following sections.

### Experimental evidence for plasmid fusion

To directly test the mechanisms driving plasmid fusion, we selected naturally occurring plasmids that carry at least one IS*257* element and belong to replicon families showing strong bioinformatic signatures of fusion (Fig. S1A–S2A). pGO1 is a conjugative plasmid from the Rep15 family ^34^ for which smaller conjugative (p.EvoF) and non-conjugative (p.Evo) derivatives have been described (Fig. S3A). ^11^ pWBG731 (p.731) is a conjugative plasmid carrying two functional replication genes, Rep20 and Rep24A, ^20^ along with a smaller non-conjugative derivative (p.144) retaining only Rep24A. As a representative small, non-transmissible plasmid, we used pUR3912 (p.3912), which belongs to the Rep13 family. ^35^

To determine whether plasmid fusion can occur at the population level, we introduced pairs of large conjugative and small non-transmissible plasmids into the same *S. aureus* strain (pGO1 or p.731 together with p.3912), allowed cultures to propagate, and extracted plasmid DNA. We designed PCR primers to specifically detect fusion junctions generated by homologous recombination between IS257 elements (Fig. S3A). Using this approach, we amplified the expected fusion products and confirmed their junction sequences by Sanger sequencing (Fig. S3B).

These results demonstrate that plasmid fusion can occur naturally through homologous recombination at IS*257* sites. However, while PCR detection establishes the occurrence of fusion events, it does not inform on their frequency within the population or the selective forces that promote their persistence, questions that we address in the following experiments.

### Antibiotic selection enriches fused plasmids

Plasmid fusion can occur naturally, but such events are intrinsically rare, in part because homologous recombination rate is low in *S. aureus*. ^36^ This rarity contrasts sharply with the high prevalence of multireplicon plasmids observed in genomic databases, suggesting that fused plasmids must be strongly selected in nature to reach detectable frequencies.

We hypothesised that strong antibiotic pressure could provide such selection, by favouring fusion between a conjugative plasmid and a non-transmissible plasmid and thereby rescuing bacterial populations under conditions that neither plasmid alone could withstand. To test this, we designed an experimental system in which a donor strain carrying both a large conjugative plasmid (pGO1, p.EvoF, or p.731) and a smaller non-transmissible plasmid (p.Evo, p.144, or p.3912) was co-cultured with a recipient strain bearing a distinct antibiotic resistance marker. Under these conditions, survival under three-antibiotic selection could only occur through plasmid fusion followed by conjugative transfer (Fig. 3A).

**Figure 3:**
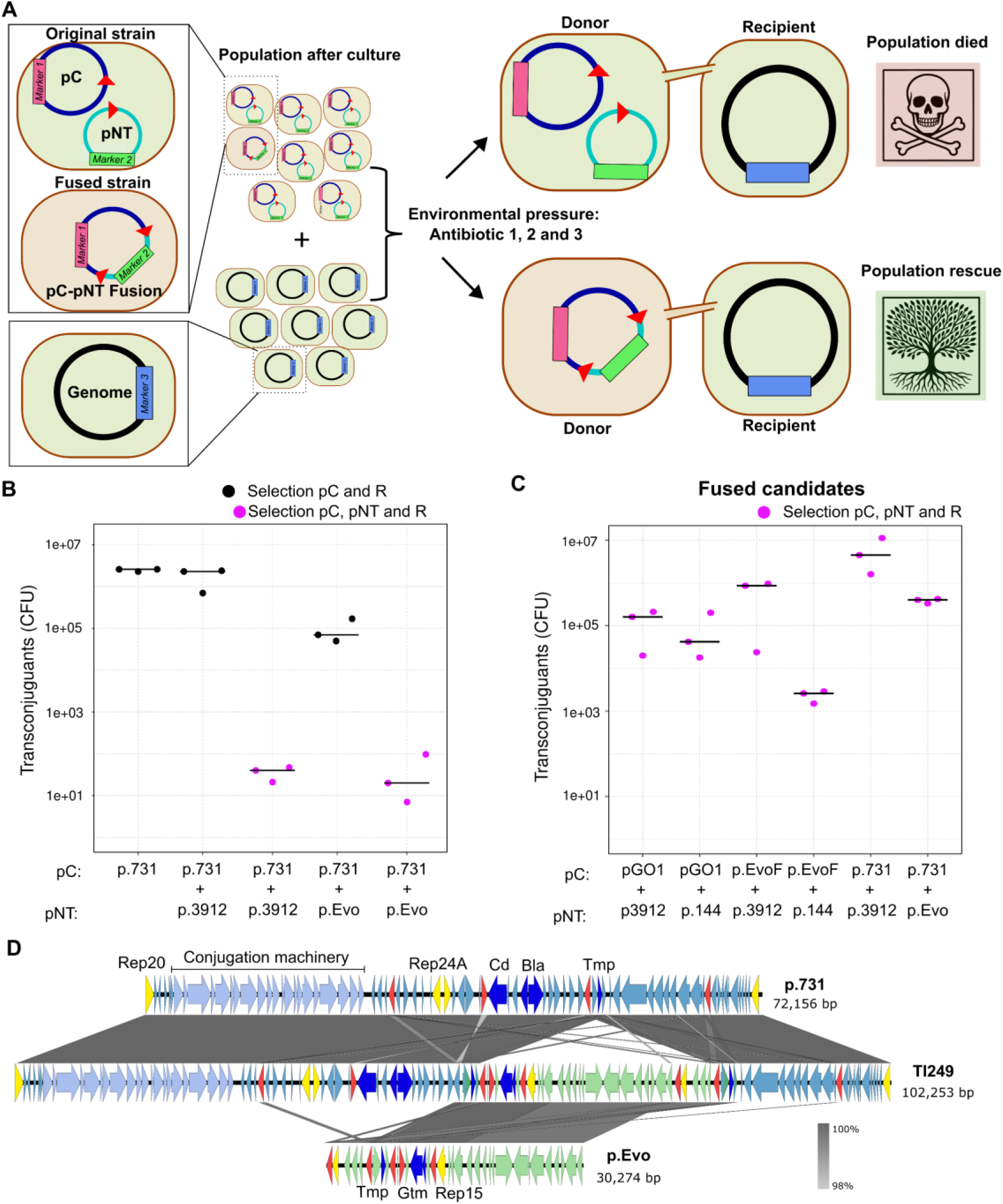
Plasmid fusion is selected during strong antibiotic pressure and rescue *S. aureus* population. Gtm: gentamycin resistance, Cd: Cadmium resistance, Tmp: Trimethoprim resistance and Bla: beta-lactam resistance. **A**-Graphical representation where plasmid fusion event would rescue the population during environmental pressure. Red Triangle represent any sequence that would bring homology for fusion between two plasmids. In the case of pGO1, p.731, p.3912, this is *IS257* gene. **B-**Experience of plasmid fusion with the conjugative pWBG731 plasmid. Conjugation controls have been used by selecting only for the conjugative plasmid and the recipient strain (Black dots). **C-** Experience of conjugation made from colony obtained after fusion experiment (in B) into a new recipient strain. **D-**Examples of plasmid fusion events sequenced by Long-Read sequencing.

Cultures were plated under two selection regimes: two antibiotics, to measure baseline conjugation efficiency, and three antibiotics, to specifically select for fused plasmids. Under two-antibiotic conditions, conjugation efficiencies were high, yielding 10^4^-10^6^ transconjugants per mL for pGO1 and p.731 (Fig. 3B, S3C). In contrast, only a small number of colonies were recovered under three-antibiotic selection. To determine whether these colonies carried fused plasmids, three isolates from each condition were used as donors in secondary conjugation assays into a fresh recipient strain. In all cases, conjugation efficiencies returned to levels comparable to those of the parental conjugative plasmids (10^3^-10^6^ transconjugants per mL; Fig. 3C), suggesting that the observed low frequencies were due to low fusion rates.

Long-read sequencing of these isolates confirmed that they carried fused plasmids containing both the conjugative and non-transmissible elements (Fig. 3D, S3D). Because fusion events occur prior to antibiotic selection (Fig. S3B), the reduction in transfer efficiency between two- and three-antibiotic conditions provides an estimate of fusion frequency in the population. Across all tested combinations, fusion events were 3-4 orders of magnitude less frequent than standard conjugation, confirming that plasmid fusion is rare under baseline conditions (Fig. 3B). Short-read sequencing of plasmids before and after fusion showed that Rep13, which allows a high copy number, is ultimately detected at a level similar to that of the large conjugative plasmid after fusion (Table S6).

Together, these results demonstrate that although plasmid fusion occurs infrequently, strong antibiotic pressure can selectively enrich fused plasmids. By conferring conjugative mobility to previously non-transmissible elements, fusion enables population survival under multidrug stress and provides a selective route by which rare fusion events can rise to prominence. This selective enrichment offers a mechanistic explanation for the widespread prevalence of multireplicon plasmids observed in genomic databases.

### Homologous recombination and transposition shape plasmid fusion

Plasmid fusion can arise through multiple molecular mechanisms, including homologous recombination (HR) and transposition, both of which can contribute to plasmid diversification. Because transposition creates large repeats that can favour HR, IS can result in both HR and transposition. To define the role of IS*257* in mediating fusion, we analysed long-read sequencing data from fused plasmids isolated during the antibiotic selection experiments (Fig. 3D; Fig. S3). In all sequenced clones, IS*257* elements flanked the fusion junctions, indicating that fusion occurred either through HR between IS*257* copies or via IS*257*-mediated transposition.

To distinguish between these mechanisms, we examined fusion junctions for signatures of transposition, specifically the presence of 7-bp direct repeats characteristic of IS*6*-family transposase activity. ^25^ In two clones (TI226 and TI228), fusion junctions were flanked by newly generated IS*257* copies and 7-bp direct repeats (Fig. 3D; Fig. S3D), consistent with transposition events. These insertions occurred at multiple sites within the small non-transmissible plasmid p.3912 and were independent of its native IS*257* element (Fig. S4C). In contrast, four clones (TI564, TI243, TI246, and TI249) lacked direct repeats, and fusion occurred directly between pre-existing IS*257* elements on each plasmid, consistent with HR-mediated co-integration (Fig. S3D).

To experimentally test whether both HR and transposition can independently mediate plasmid fusion, we constructed derivatives of p.3912 lacking IS*257* (p.noIS), as well as a variant containing an alternative region of homology (the trimethoprim resistance gene) shared with p.EvoF and p.731 (Fig. S4A). Under selective conditions favouring fusion, all plasmid variants supported population rescue following conjugation with p.731 and p.EvoF (Fig. S4B). Long-read sequencing confirmed plasmid fusion in all cases and revealed junction architectures consistent with either HR or transposition (Fig. S4C–S4F). Analysis of transposition-derived junctions identified a conserved 7-bp direct repeat motif, allowing definition of a consensus insertion site (TTTWTAD) (Fig. S4C–S4D).

Together, these results demonstrate that plasmid fusion can be mediated by both homologous recombination and transposition. Moreover, short regions of sequence homology, including conserved replication genes, resistance determinants, or transposase sequences, are sufficient to promote co-integration. This mechanistic flexibility provides a basis for the widespread generation of plasmid diversity and illustrates how mobile genetic elements facilitate rapid adaptation under selective pressures such as antibiotic exposure.

### Plasmid size does not impose a fitness cost after fusion

Although plasmid fusion is frequent and widely represented in genomic databases, very few *Staphylococcus* plasmids exceed 45 kb, even among those with clear fusion signatures (Fig. 1B). Notably, 45kb corresponds to the maximum packaging capacity of most *Staphylococcus* transducing phages, ^10^ supporting the idea that phage-mediated transfer imposes an upper limit on plasmid size. ^11^ We considered three non-mutually exclusive explanations for this pattern: (i) a fitness cost associated with carrying large plasmids, (ii) resolution of fused plasmids after co-integration, and (iii) selection for deletions, potentially driven by phage-mediated transfer constraints.

To test whether increased plasmid size or the presence of multiple replication systems imposes a fitness cost, we compared the growth of bacteria carrying fused plasmids with that of strains carrying their unfused parental plasmids. Using fused plasmids generated in our experimental system, we first assessed whether plasmid size correlated with reduced growth (Fig. 4A). No consistent size-dependent fitness cost was detected. For example, the 76-kb fused plasmid derived from pGO1 grew comparably to its parental plasmids pGO1 (54 kb) and p.EvoF (42 kb). The smallest derivative, p.Evo, displayed slightly faster growth, likely reflecting the absence of conjugation machinery as observed in a *K. pneumoniae* plasmid. ^37^ Similarly, in the p.731 background, fused plasmids of 78 kb and 102 kb grew as well as, or slightly better than, the smaller p.144 (26 kb) and the original p.731 plasmid which curiously showed the poorest growth.

**Figure 4:**
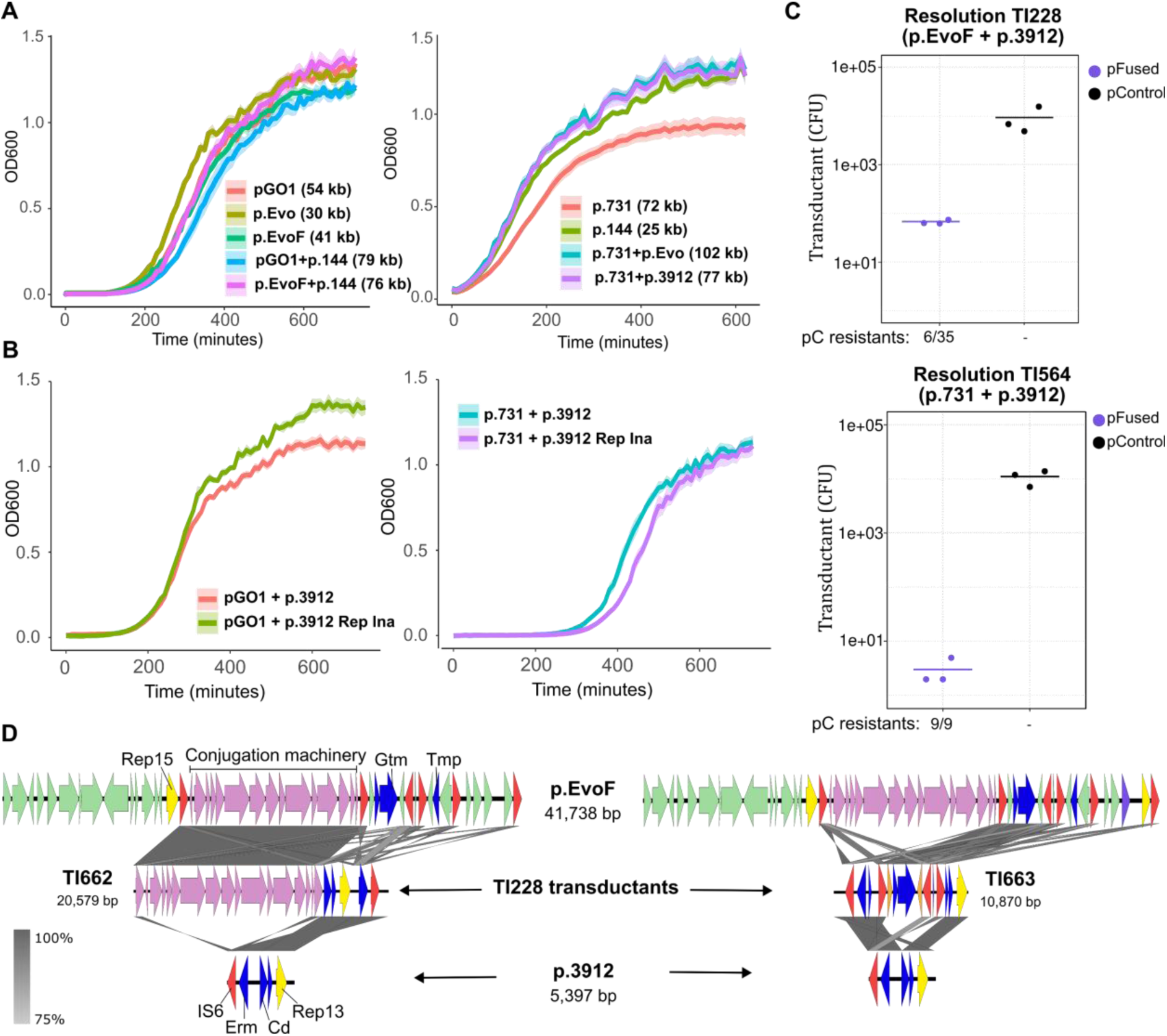
Plasmids size limitation is not linked to fitness or resolution mechanism after plasmid fusion. **A**-Growth curve of plasmid issued from pGO1 family, his fused version and the smaller versions and Growth curve of plasmid issued from p.731 family, his fused version and the smaller versions. **B**-Growth curve of plasmid having multi-replicons that are functional or not obtained after plasmid fusions. **C**-Plasmid resolution efficiency from TI228 (p.EvoF-p.3912 fusion plasmid) and TI564 (p.731-p.3912 fusion plasmid), detected after phage transduction of and selection of the small plasmid pUR3912, that should select resolved plasmid after fusion. Number of colonies still resistant to the conjugative plasmid is represented. **D-** Example of sequenced clones (TI662 and TI663) obtained after the plasmid resolution assay from TI228 (p.EvoF-p.3912 fusion plasmid).

We next examined whether carrying two active replication systems within a single plasmid affected bacterial fitness. We compared growth of strains harbouring fused plasmids in which one Rep gene was naturally inactivated during fusion with those retaining two functional Rep genes (Fig. 4B). Again, no significant differences in growth were observed.

Together, these results indicate that neither increased plasmid size nor the presence of multiple active replication systems imposes a detectable fitness cost under the conditions tested. The predominance of smaller plasmids in natural populations therefore cannot be explained by fitness limitations alone, implicating alternative selective forces, such as deletion processes or phage-mediated constraints, in maintaining compact plasmid sizes.

### Resolution and deletion diversify fused plasmids

Previous studies have shown that plasmids generated by transposition can sometimes resolve back into their original components. ^25,38,39^ To determine the intrinsic fates of fused plasmids after co-integration, we examined the outcomes of fusion events independently of their long-term selective success.

Using fused derivatives of pGO1–p.3912 and p.731–p.3912 generated in earlier experiments, we exploited phage-mediated transduction while selecting for resistance encoded by the smaller plasmid. Because large, fused plasmids exceed the DNA packaging capacity of most staphylococcal phages, colonies recovered under these conditions were expected to reveal post-fusion processing events, including resolution or deletion (Fig. 4C).

Analysis of recovered colonies revealed two distinct mechanistic outcomes. In two sequenced clones (TI267, TI273), the original small plasmid was regenerated, indicating resolution of the co-integrated plasmid back into its parental components. In parallel, a second class of colonies retained resistance markers associated with the large conjugative plasmid. Among 35 colonies examined for pGO1–p.3912 and 9 for p.731–p.3912, 6 and 9 colonies, respectively, displayed this phenotype (Fig. 4C).

Long-read sequencing showed that these latter isolates carried novel intermediate-sized plasmids generated by deletion rather than simple resolution (Fig. 4D). In one case, deletion of the pGO1 backbone resulted in a p.3912 replicon plasmid (Rep13) that contained the full pGO1 conjugation machinery, effectively converting the original non-transmissible Rep13 plasmid sub-family into a conjugative plasmid. In all sequenced cases, deletion junctions were consistent with homologous recombination between IS257 elements (Fig. 4D).

Deletion events were also detected in the absence of phage infection. Using plasmids carrying multiple IS257 copies, we amplified and sequenced recombination scars consistent with IS257-mediated deletions occurring naturally at the population level (Fig. S5).

Together, these results demonstrate that fused plasmids act as diversification intermediates, from which multiple evolutionary outcomes can arise, including resolution back to parental plasmids and deletion-driven formation of novel chimeric elements.

### Phage pressure selectively enriches small plasmid derivatives

While fused plasmids generate multiple derivative forms, the relative abundance of these outcomes in natural populations is expected to depend on ecological selection. In particular, bacteriophages impose strong physical constraints on DNA packaging, potentially favouring smaller plasmid variants capable of being transduced.

To test whether phage pressure selectively enriches specific post-fusion outcomes, strains carrying multireplicon plasmids larger than 45 kb were infected with phages 80α or □2339 and transduction experiment were performed into a recipient strain under defined selection regimes. We predicted that phage-mediated transfer would act as a filter, enriching for plasmid variants compatible with phage packaging carrying various combinations of genes as observed previously (Fig. 4D).

Consistent with this prediction, transduction recovered a diverse spectrum of plasmid derivatives in all plasmid–phage combinations tested (Fig. 5A-B). Long-read sequencing revealed plasmids of intermediate sizes (33.5-44.5 kb) with distinct architectures, including deletion derivatives, resolved parental plasmids, and novel combinations of resistance genes and conjugation modules (Fig. 5; Fig. S6).

**Figure 5:**
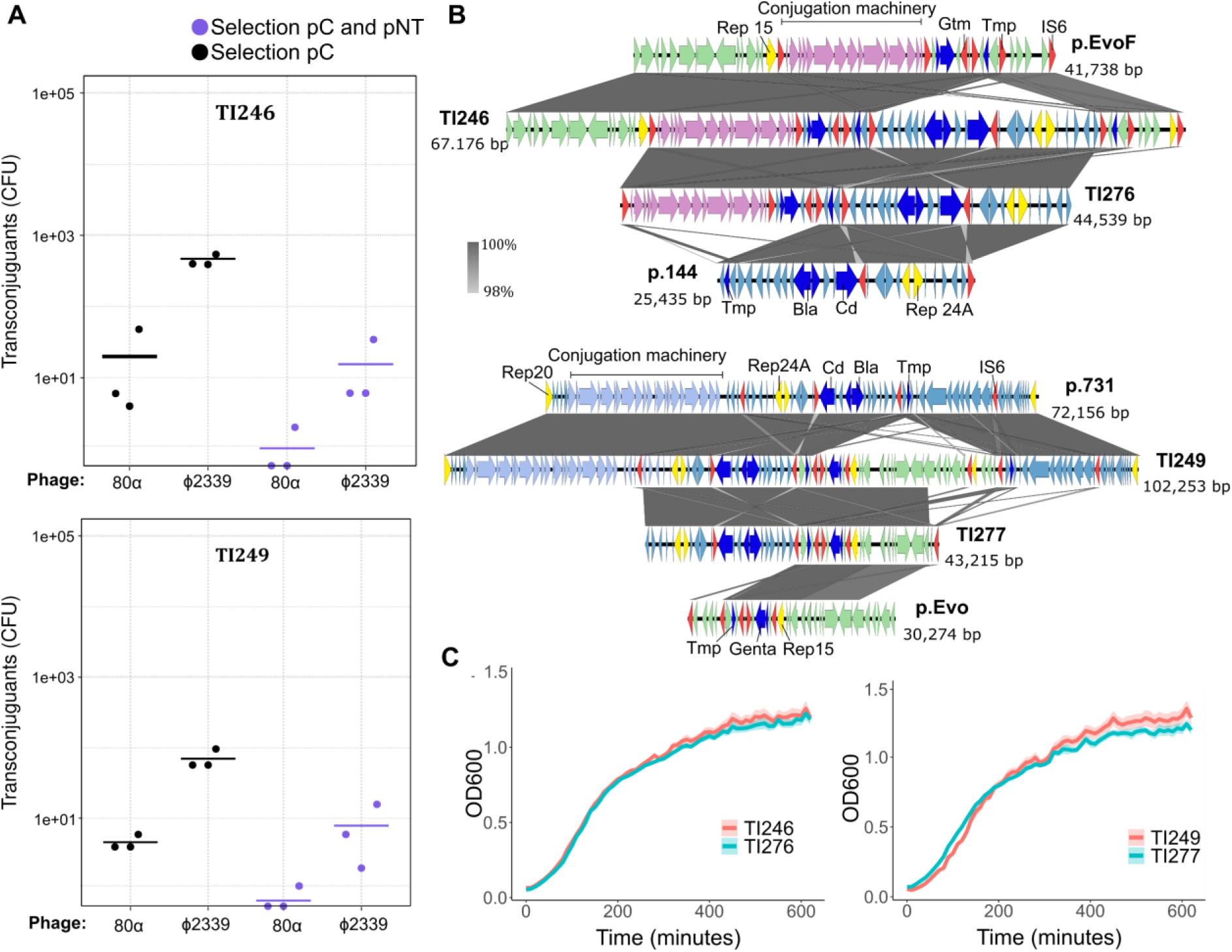
Phage select for diverse intermediate size plasmid derivate. **A**-Transduction of TI246, the fused version p.EvoF/p.144 and TI249, the fused version p.731/p.Evo. **B**-Sequence alignment of derivates obtains after transduction. **C-** Growth curve of plasmid fused version vs transduce version.

Importantly, growth assays showed no detectable fitness cost associated with these smaller derivatives relative to their fused precursors (Fig. S6), indicating that their enrichment was driven by transfer efficiency rather than growth advantage.

Together, these results demonstrate that bacteriophage pressure does not generate plasmid diversity de novo, but selectively amplifies specific outcomes produced by plasmid fusion, favouring smaller, transducible variants. This selective filtering explains how plasmid size remains constrained in nature while preserving extensive genetic diversity.

## Discussion

Plasmids are central engines of bacterial adaptation, yet the evolutionary forces shaping their diversity remain poorly understood. Integrating our bioinformatic, genetic, and experimental data, we identify a plasmid life cycle that operates at the population level and is driven by mobile genetic elements. In *S. aureus*, this life cycle is characterised by alternating phases of fusion, diversification, and ecological selection (Fig. 6). Under antibiotic pressure, rare fusion events generate multireplicon plasmids that transiently expand genetic repertoires and mobilize previously non-transmissible plasmids, rescuing bacterial populations under strong antibiotic stress. These fused plasmids do not represent stable evolutionary endpoints but instead act as transient diversification intermediates. They can resolve back into parental components or undergo deletion to form intermediate-sized chimeric plasmids that retain key functional modules. Phage-mediated transduction then selectively enriches smaller, transducible derivatives, imposing a strong constraint on plasmid size while preserving genetic diversity. Together, this fusion-diversification-selection process explains how plasmid populations maintain conserved backbones, constrained size distributions, and extensive accessory gene diversity without preserving strict plasmid identity.

**Figure 6:**
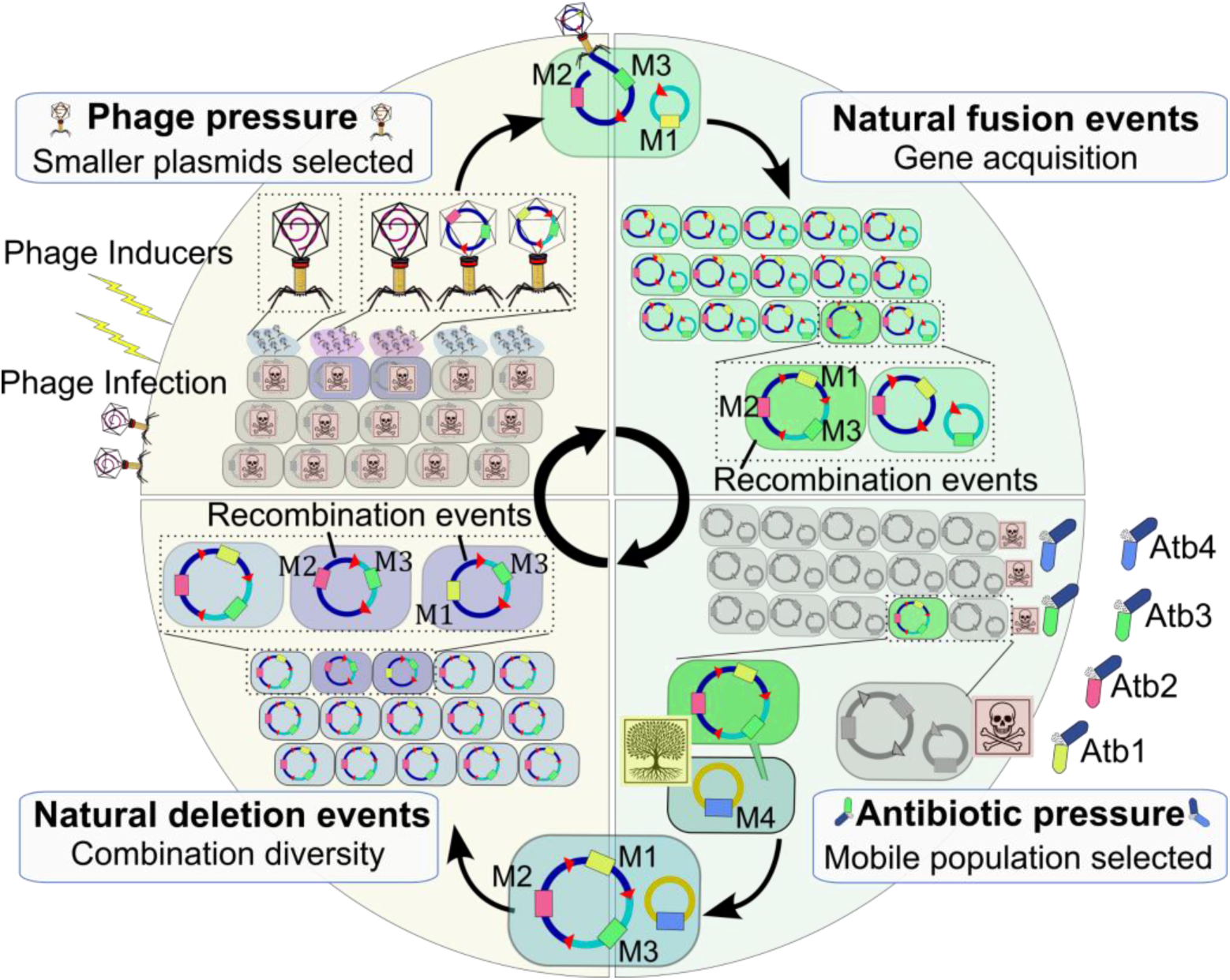
Model of plasmid evolution in *Staphylococcus.* Rare fusion events, mediated by homologous recombination or transposition, combine distinct plasmids into single multi-replicon elements, expanding gene content and transfer potential. Antibiotic pressure selectively enriches these fused plasmids, rescuing populations under stress. Rare deletion events retain essential genes and transmissibility and create new combinations of genes. Phage predation imposes size constraints by killing the whole population and selecting plasmids that can fit into their capsid and be transferred to another cell.

Insertion sequences (ISs) emerge as central catalysts of plasmid diversification. They facilitate both fusion and deletion through homologous recombination and transposition, enabling rapid structural rearrangements across short regions of homology. This role is consistent with observations in other bacterial species, where IS6-family elements mediate plasmid co-integration via similar mechanisms. ^26,30,40^ Importantly, IS-mediated rearrangements do not merely generate transient intermediates but can give rise to stable plasmid variants that persist under selection. Conjugative plasmids display particularly high fusion frequencies, mirroring patterns reported in *Escherichia coli* and other species where IS density is elevated near antimicrobial resistance genes. ^24,41,42^ Beyond IS elements, other conserved genetic features, including resistance cassettes (Fig. S4) and replication genes, can also serve as substrates for homologous recombination. While replication genes represent potential recombination targets, plasmid incompatibility limits the stable coexistence of identical replicons within the same cell, ^43^ suggesting that plasmid fusion is shaped by a balance between recombination potential and incompatibility constraints.

Although multireplicon plasmids have been reported for decades, they were long regarded as rare or exceptional products of plasmid evolution, often described as isolated curiosities rather than as evolutionarily consequential entities. Recent population-scale analyses, particularly in *Enterobacteriaceae*, have challenged this view by revealing a historical expansion of multireplicon plasmids during the antibiotic era, suggesting that they may play a major role in contemporary resistance evolution. ^22,23^ However, these studies did not determine precisely how frequently plasmid fusion occurs, how such rare events become evolutionarily significant, or whether multireplicon plasmids represent stable endpoints or transient stages within broader evolutionary dynamics. Our results directly address these gaps by showing that multireplicon plasmids are common in *Staphylococcus*, arising repeatedly through MGE-driven fusion that produces short-lived large plasmids functioning as key intermediates in a dynamic evolutionary process. Plasmid fusion generates large multireplicon plasmids that enhance genetic content and provide plasmid mobility. In *Staphylococcus*, multireplicon plasmids are widespread in genomic databases, representing nearly half of all plasmids analysed but we showed that large plasmids are rare in nature (6% of *Staphylococcus* plasmids > 45kb). This apparent paradox is resolved by the transient nature of fusion events and the branching evolutionary trajectories that follow (Fig 3, 6). After fusion, plasmids can resolve into their original components at low frequency or undergo deletion to generate novel chimeric plasmids retaining key functional modules. These branching outcomes substantially increase accessory gene diversity within bacterial populations, generating a spectrum of plasmid architectures from a single fusion event.

Environmental pressures then determine which of these variants persist. Fitness cost has often been invoked as the primary driver of plasmid deletion. ^44–47^ However, we show that plasmid size alone does not impose a clear fitness cost in *S. aureus*. Instead, bacteriophage predation imposes a strong physical constraint on plasmid size by selectively enriching variants compatible with phage-mediated transduction. Large fused plasmids are unable to be fully packaged and are lost, whereas deletion derivatives and resolved plasmids that fall below phage packaging limits are preferentially propagated. This size-dependent filtering is further reinforced by staphylococcal pathogenicity islands (SaPIs), which package only small DNA molecules, thereby favouring even more compact plasmid architectures. ^11^

Altogether, our results provide mechanistic insight into plasmid evolution: antibiotics select for rare fusion events that combine multiple resistance determinants, while phage and satellite pressure constrain plasmid size by selecting deletion-derived variants. A defining feature of the plasmid life cycle described here is that it relaxes, rather than enforces, the conservation of genetic architecture across generations. In contrast to many MGEs, whose life cycles tend to preserve a core architecture, it prioritizes diversification over replication fidelity. Each fusion event seeds multiple evolutionary trajectories, producing a branching set of descendants rather than a single reverting form. This controlled diversification is not a limitation of the system but its principal evolutionary advantage, allowing plasmid populations to explore a wider genetic space while remaining constrained by transmission ecology. We propose that this highly diversifying, MGE-driven life cycle represents a general principle of plasmid evolution, particularly in species where multiple mobile elements coexist and interact. This process shapes the mosaic architecture of modern resistance plasmids and aligns with recent work showing that population-level diversity enhances survival under fluctuating antibiotic and phage pressures. ^48^

An important layer not explicitly incorporated into our model is the role of plasmid-encoded phage defence systems. Defence mechanisms such as restriction–modification systems or abortive infection modules may influence which plasmid variants persist following fusion or deletion. Large conjugative plasmids combining antimicrobial resistance with effective phage defence could be preferentially retained. Conversely, smaller plasmids may escape restriction barriers more efficiently ^49,50^ and achieve higher resistance expression due to increased copy number. ^51^ Incorporating phage defense systems into future models will further refine how plasmid mobility, gene content, and architecture are optimized under complex ecological pressures.

Although this study focuses on *S. aureus*, the underlying principles are likely generalisable. Multireplicon plasmids, IS-mediated fusion, and deletion events are reported in Enterobacteriaceae, *Klebsiella*, *Salmonella*, *Acinetobacter*, and *Citrobacter*, ^13,16–19,23,40,52^ often with major consequences for plasmid mobility and resistance dissemination. ^53^ Plasmids may therefore represent an extreme case of evolvable mobile elements, in which repeated interactions with other MGEs amplify genetic diversity beyond what is observed in more canonical life cycles.

We conclude that plasmid diversification arises from a defined evolutionary life cycle driven by mobile genetic elements and ecological selection. By framing plasmid evolution as a cyclical, population-level process rather than the persistence of individual elements, our model explains the rapid emergence, stability, and global dissemination of multidrug-resistant plasmids. This conceptualization provides a framework for anticipating resistance evolution and highlights the need to consider phage-plasmid interactions when designing strategies such as phage therapy to combat antimicrobial resistance.

## Materials and Methods

### Oligonucleotides and plasmids

All oligonucleotides used in this study were supplied by Sigma and are listed in Table S5-A. All plasmids constructed and used in this study are listed in Table S5-B.

### Bacterial strains and growth conditions

The bacterial strains used in this study are listed in Table S5-C. For all experiments, *S. aureus* was grown in Tryptic Soy Broth (TSB) at 37 °C with shaking (180 rpm) overnight until stationary phase. For enumeration of colony-forming units (CFU) on solid media, cultures were plated on Tryptic Soy Agar (TSA) and incubated at 37 °C. When required, the media was supplemented with the following antibiotic concentrations: gentamicin 10 µg/ml; erythromycin 10 µg/ml; trimethoprim 20 µg/ml; chloramphenicol 10 µg/ml; CdCl₂ 100 µM; tetracycline 3 µg/ml; rifampicin 10 µg/ml.

When using *Saccharomyces cerevisiae (*provided by Dr. Carole Lartigue, INRAE) as a cloning platform, *S. cerevisiae* VL6-48N was grown in YPDA broth (Takara, 630464) overnight at 30 °C with shaking (200 rpm). When required, minimal SD medium (SD-Trp broth; Takara, 630411 and 630413) was used at 30 °C with shaking at 220 rpm.

### Bioinformatic analyses

3,076 *Staphylococcus* plasmids were downloaded from PlasmidScope database ^54^ (version 2024-12-04, all the plasmids were collected from PLSDB ^55^). After removing potential chromosomal contigs and merging identical sequences, 2,564 non-redundant complete plasmids in a size range of 2-170 kb were kept for the further analysis (Supplementary Table S1).

Plasmid mobility typing was performed using MacSyFinder v2.1.4 ^56^ with CONJScan ^57^ model v2.0.1. OriT sequences were searched using BLASTn ^58^ (v 2.17.0 with parameter - task blastn-short, identity > 80% and coverage > 80%) with 117 experimentally valid oriT sequences from OriTDB v2.0, ^59^ the best hit from overlapping hits was kept. Plasmid mobility types were classfied into plasmid encoding oriT sequence (pOriT), plasmid encoding relaxase (pMOB), conjugative plasmid (pCONJ), decayed conjugative plasmid (pdCONJ), and plasmid lacking mobility (pNT). 23 plasmids with mobility type “pdCONJ” are grouped into “pMOB” in the further analysis.

Plasmid replicon typing was performed using PlasmidFinder v2.1.1 ^60^ and replicon database v2.2.0 (with parameters -t 0.8 -l 0.6, identity > 80% and coverage > 60%), the best hit from overlapping hits was kept (Supplementary Table S2). Replicon co-occurrence networks were plotted using Cytoscape v3.10.3. ^61^

Insertion sequence IS*6* and IS*Bli29* were detected using digIS v1.2 ^62^ (Supplementary Table S3) and IS*6* transposase sequences were cross valided using Blastn with a pre-build blast database of Tncentral ^63^ and ISFinder ^64^ (version 2025-05-16), for each query the best hit was kept (Supplementary Table S4).

### Detection deletion and fusion event in *Staphylococcus*

Bacterial strain *S. aureus* RN4220 containing one or combinations of plasmids was cultured overnight, and plasmids were extracted from 1 mL of culture using the ZR BAC DNA Miniprep Kit (Zymo Research, D4049). PCRs were performed using Advantage II HF polymerase (Takara), following the manufacturer’s recommendations.

### Big plasmid construction

The natural plasmid pGO1 lacking the conjugative machinery and containing a chloramphenicol resistance gene was constructed in the non-commercial *Saccharomyces cerevisiae* VL6-48N strain, as previously described ^65,66^ and ^67^ with several modifications. Overlapping PCR products and YAC (200 ng of each PCR product and 100 ng of linearized YAC in 10 µL water) were co-transformed into VL6-48N. Individual yeast colonies were picked and streaked onto SD-Trp plates, incubated for 2 days at 30°C, and then patched onto fresh SD-Trp plates and incubated for another 2 days at 30°C. Total genomic DNA was extracted from yeast transformants according ^65^. PCRs were performed using Advantage II HF polymerase (Takara), following the manufacturer’s recommendations.

### Small plasmid construction

Small p.3912 derivatives lacking IS256, or in which IS256 was replaced by a sequence allowing homologous recombination with pGO1 and p.731, and carrying the trimethoprim resistance gene, were constructed in *S. aureus* RN4220. Overlapping PCR fragments were generated and assembled using the HiFi DNA Assembly Kit (NEB). The reaction products were dialyzed on a membrane filter for 5 min, and the remaining mixture was transformed into *S. aureus* RN4220 (2,200 kV, 100 Ω, 2 mm) and plated on TSA containing erythromycin. Colonies were then analyzed, and plasmids were sequenced.

### Bacterial growth curves

Growth kinetics were assessed using *Staphylococcus aureus* RN4220 derivatives carrying the indicated plasmids. Overnight cultures were inoculated into fresh Tryptic Soy Broth (TSB) at a 1:100 dilution and grown at 37 °C with shaking (180 rpm). Cultures were transferred to 96-well flat-bottom plates (200 μL per well; three technical replicates per strain), and optical density at 600 nm (OD600) was measured every 10 min for 24 h using a microplate reader with continuous orbital shaking between reads.

### Phage infection

Bacterial cultures containing the appropriate plasmid were grown to an OD600 of 0.15. One milliliter of culture was mixed with 100 μL of phages supplemented with 5 mM CaCl₂ and incubated at 30°C for 3 h, followed by overnight incubation at room temperature. The resulting phage lysate was filtered to remove remaining bacterial cells.

### Phage titer

*S. aureus* RN4220 was grown overnight, then subcultured at a 1:50 dilution into fresh TSB and grown to an OD600 of 0.2. Bacterial lawns were prepared by mixing 100 μL of cells with phage top agar (PTA, supplemented with CaCl₂ to a final concentration of 5 mM) and pouring the mixture onto PBA plates supplemented with 5 mM CaCl₂. Serial dilutions of phage lysates were prepared in phage buffer (50 mM Tris, pH 8; 1 mM MgSO₄; 4 mM CaCl₂; 100 mM NaCl) and spotted onto the bacterial lawns, which were then incubated overnight at 37°C. Plaques were counted and used to quantify the phage produced during infection.

### Plasmid transduction

*S. aureus* RN4220 was grown in TSB to an OD600 of 1.4. For transduction experiments, 100 μL of phage lysates were added to each culture along with 5 mM CaCl₂. The cultures were incubated statically at 37°C for 30 min and then plated onto TSA plates containing 17 mM sodium citrate and the appropriate antibiotic for selection of the expected plasmid. Plates were incubated for 24–48 h, and colonies were counted. The transfer efficiency (TE) per donor cell was calculated as the number of Transductant Units (TrU) per mL divided by the number of donor colony-forming units (CFU) per mL (6.5 × 10□ CFU/mL for a culture with an OD600 of 0.15).

### Statistical analyses

The enrichment of different replicon family in multiple- / single-replicon plasmid was tested using Fisher’s exact test in contingency analysis and correction for multiple comparisons with false discover rate (FDR) ^68^. Plasmid encoding multiple- / single-replicon was modeled using a binomial generalized linear model using glm function (R package stats v4.4.3, with parameter family = binomial (link=’logit’)). The effect of plasmid size (log10 transformed), IS_presence (plasmid encoding transposase of IS*6*/ IS*Bli29* or not), and plasmid mobility (mobile (i.e. pOriT, pMOB, pdCONJ, pCONJ) or non-mobile (pNT)) are tested. All statistical analysis were conducted in R v4.4.3.

## Supporting information

Table S1. *Staphylococcus* plasmids analysis.

Table S2. *Staphylococcus* plasmids replicon classification.

Table S3. *Staphylococcus* plasmids IS detection.

Table S4. *Staphylococcus* plasmids IS257 detections.

Table S5. Strains and oligonucleotides used in this study.

Table S6: Plasmid copy numbers analysis.

## Acknowledgements and funding sources

This work was supported by grants MR/X020223/1, MR/M003876/1, MR/V000772/1 and MR/S00940X/1 from the Medical Research Council (UK), BB/V002376/1 and BB/V009583/1 from the Biotechnology and Biological Sciences Research Council (BBSRC, UK), and EP/X026671/1 from the Engineering and Physical Sciences Research Council (EPSRC, UK) to J.R.P. Thank you to Dr Carole Lartigue from INRAE for providing the VL6-48N strain, to Dr Joshua Ramsay from Curtin University for providing pWBG731 strain and to Daniel Sin for help with plotting.

## Author contributions

J.R.P. and T.I. conceived the study; T.I and Y.W. conducted the experiments; T.I., Y.W., E.R., and J.R.P. analyzed the data; J.R.P. and T.I. wrote the manuscript.

---

**Figure S1:**
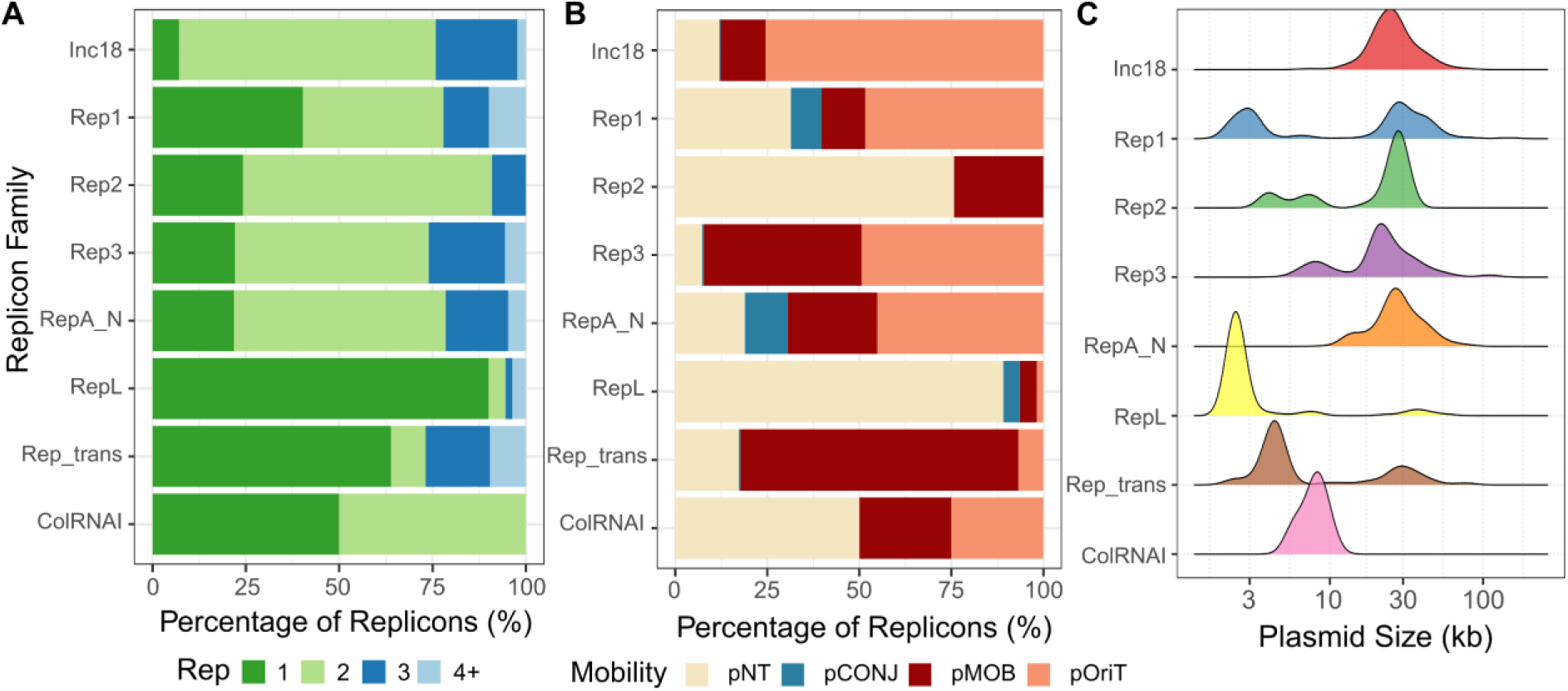
Distribution of plasmid replicon families in *Staphylococcus*. **A-** Proportion of plasmid encoding 1-4+ replicon(s) within each replicon family, plasmids encoding more than four replicons are merged into “4+” group. **B-** Distribution of plasmid mobility types within replicon family. **C-** Comparison of plasmid size (log10 transformed) across replicon family.

**Figure S2:**
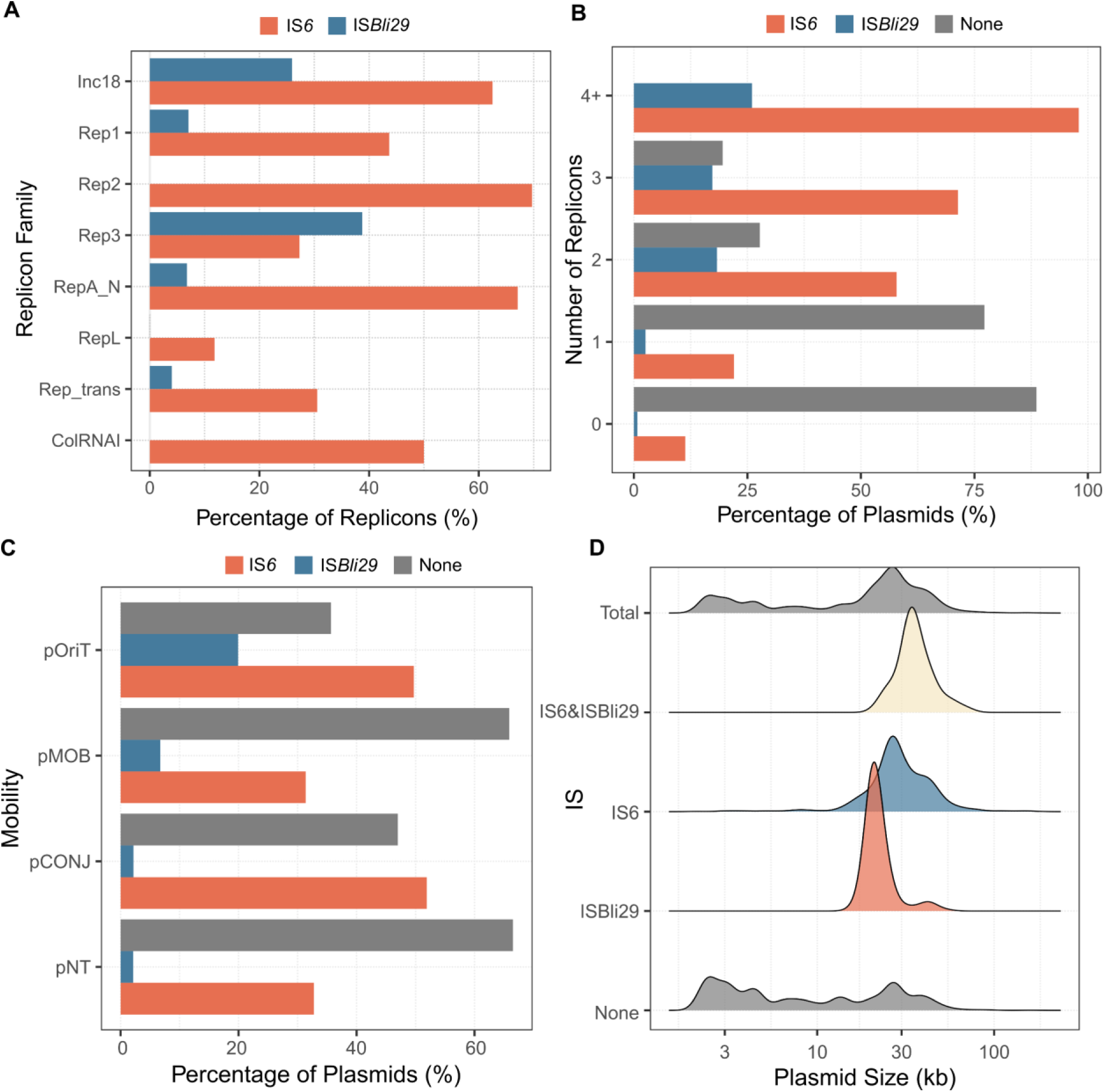
Distribution of Insertion Sequence (IS) IS*6* and IS*Bli29* in *Staphylococcus* plasmids. **A-** Frequency of IS*6* and IS*Bli29* in plasmids grouped by replicon family. **B-** Proportion of IS*6* and IS*Bli29* encoding plasmids with 1-4+ replicon(s), plasmids with more than four replicons are merged into “4+” group, plasmid encoding replicon not identifiable are in group “0”. **C-** Proportion of IS*6* and IS*Bli29* encoding plasmids by mobility types. **D-** Comparison of plasmid size by encoding IS*6*, IS*Bli29*, both, and neither. (Note: 97.6% of the IS*6* here is cross valid by ISfinder as IS*6* family clade f, i.e. IS*257*; IS*Bli29* here is the ORF4 which encodes a transposase with a Mu transposase C-terminal domain, pfam accession PF09299).

**Figure S3:**
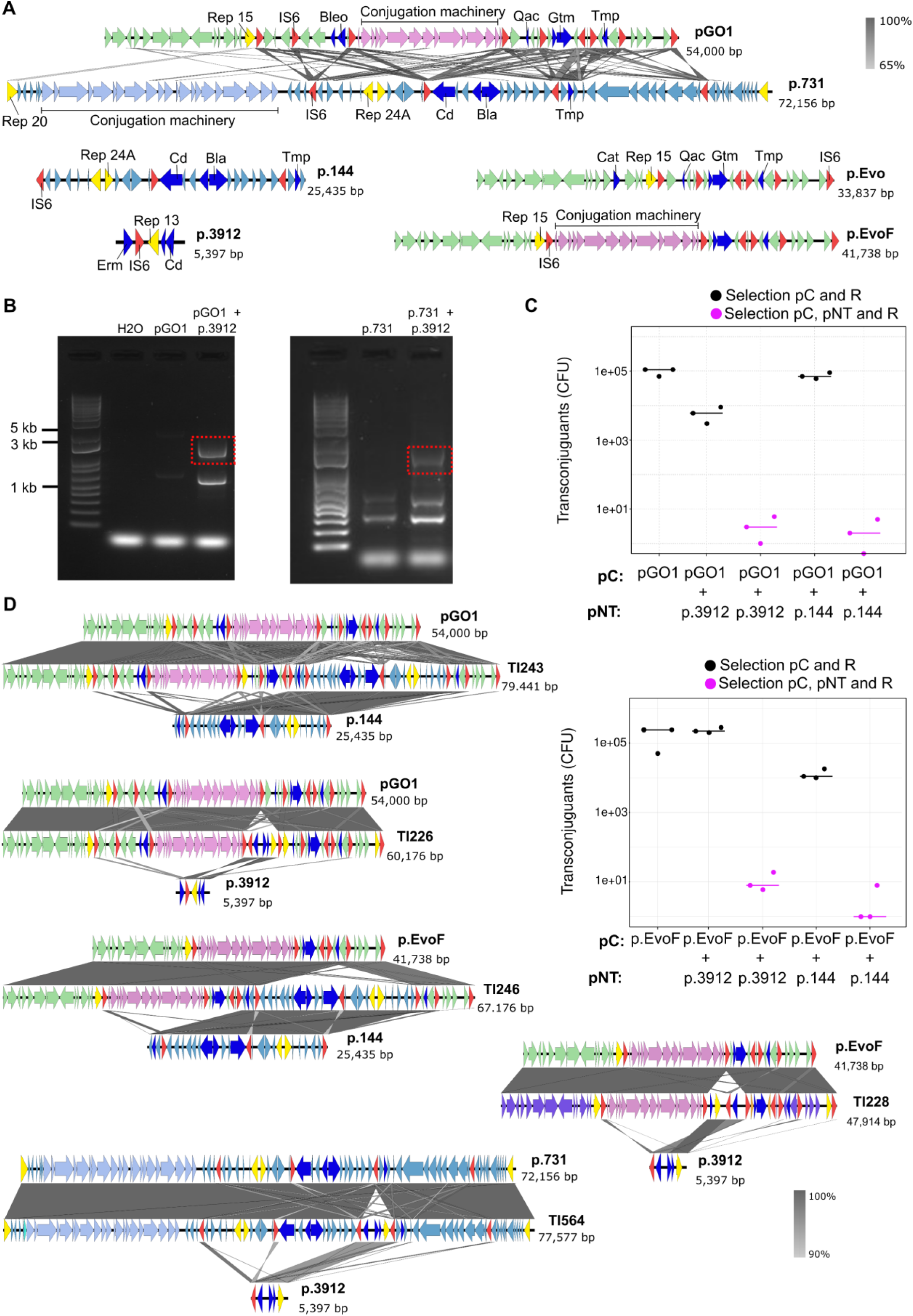
Plasmid fusion assays. pC (Conjugative plasmid), pNT (non-transmissible plasmid), R (Recipient). Gtm: gentamycin resistance, Cd: Cadmium resistance, Tmp: Trimethoprim resistance, Erm: Erythromycin resistance. Bleo: Bleomycin resistance, Bla: beta-lactam resistance and Qac: quaternary ammonium compounds resistance. **A**-Plasmids map used in this study. Primer designed for the detection of fusion events on pGO1 and p.731 plasmids with p.3912 are represented. **B**-Detection of fusion events within pGO1 and p.3912 IS257 and p.731 and p.3912. **C**-Experience of plasmid fusion with the conjugative pGO1 and p.EvoF plasmid. Conjugation controls have been used by selecting only for the conjugative plasmid and the recipient strain (Black dots). **D-** Other examples of plasmid fusion events sequenced by Long-Read sequencing.

**Figure S4:**
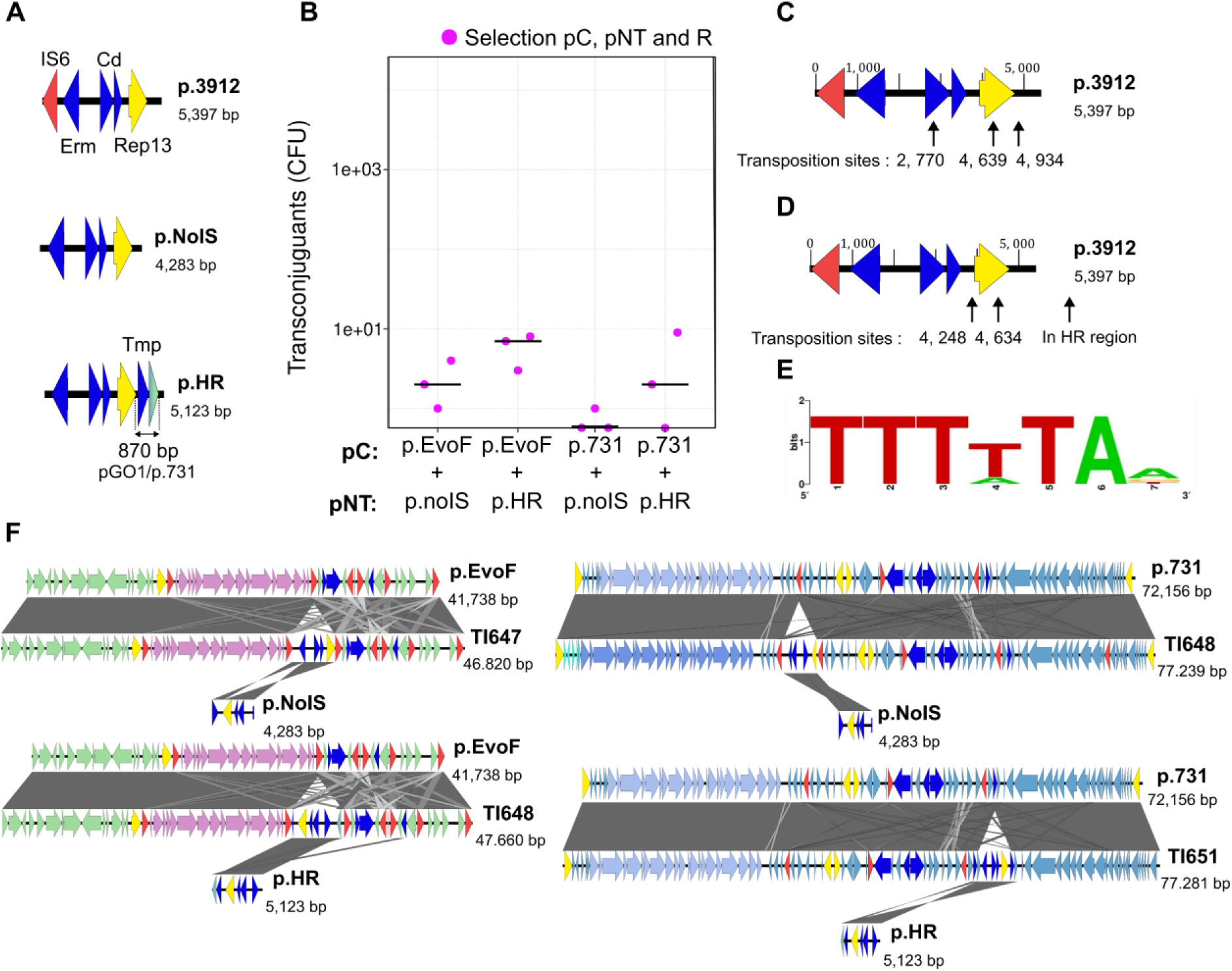
Plasmid fusion is issued from both HR or Tp events. pC (Conjugative plasmid), pNT (non-transmissible plasmid), R (Recipient). Cd: Cadmium resistance, Tmp: Trimethoprim resistance and Erm: Erythromycin resistance. **A**-Derivative plasmid from p.3912 after *IS6* gene removing or replaced by another gene find in both pGO1 and p.731. **B-** Experience of plasmid fusion with p.3912 derivates to test homologous recombination (HR) and transposition (Tp) mechanisms. **C-** Sites of transposition in pC-p.3912 fusion plasmids. **D**-Sites of transposition in pC-p.3912 derivates fusion plasmids. **E**-Consensus motif for transposition site. **F-** Examples of plasmid fusion events sequenced by Long-Read sequencing.

**Figure S5:**
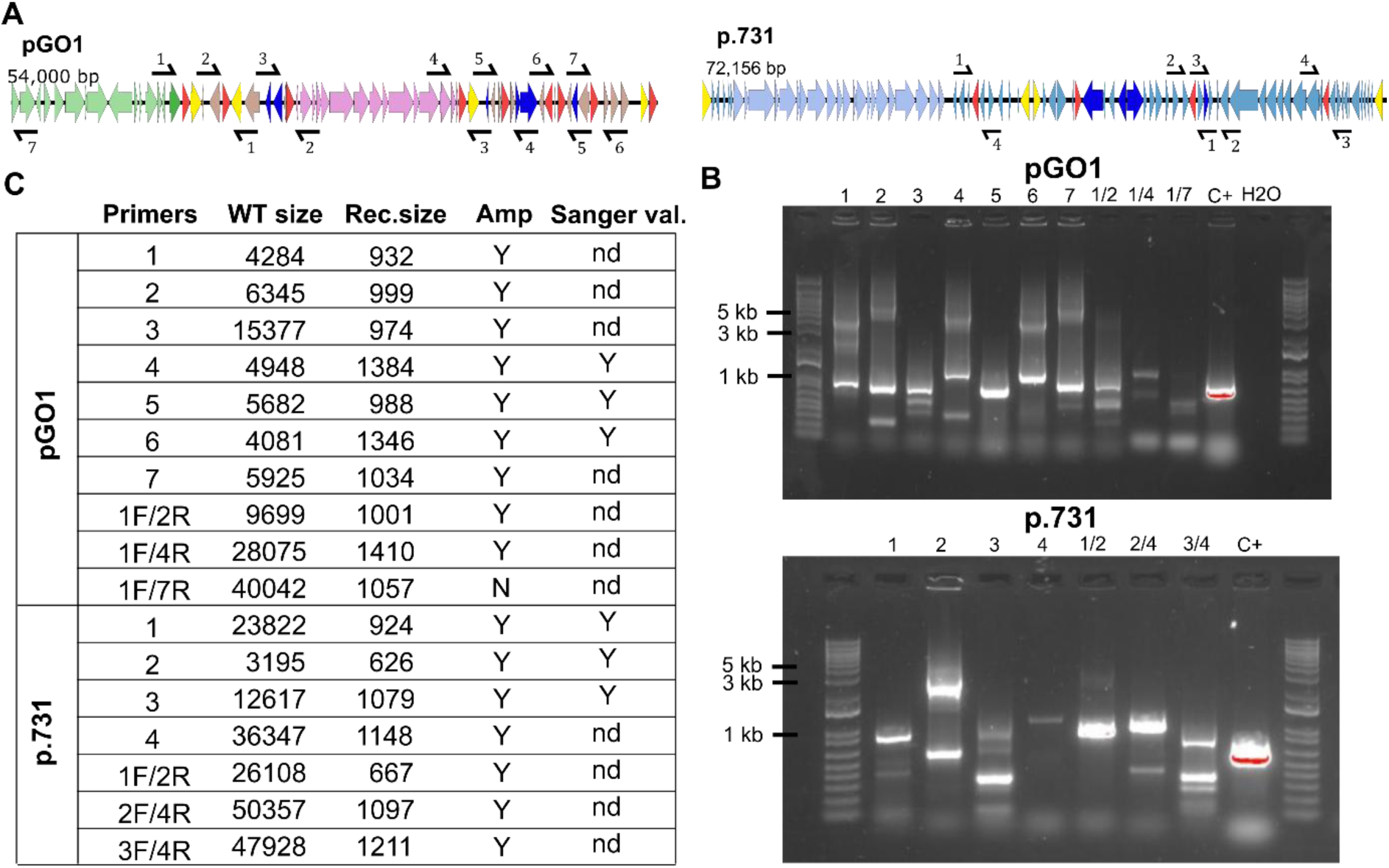
Detection of deletion events in bacterial cell population. **A**-Design of the primers for detection of deletion in pGO1 and p.731 between different copies of IS257. **B**-Detection by PCR of Recombination scars. **C**-Recapitulative table of deletion detection results.

**Figure S6:**
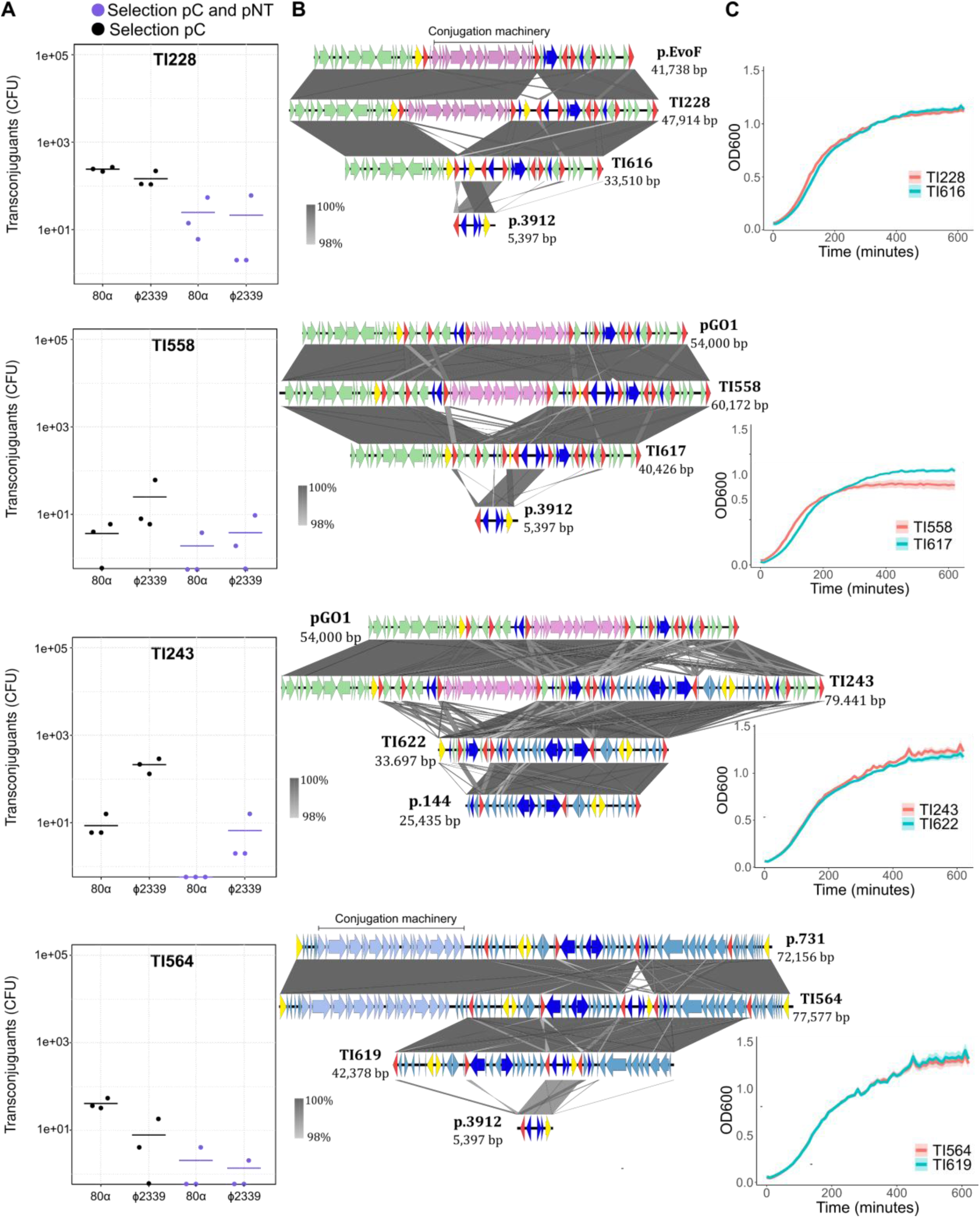
Phage select for diverse intermediate size plasmid derivate. **A**-Transduction of TI228, the fused version p.EvoF/p.3912, TI558 the fused version pGO1/p.3912, TI243 the fused version pGO1/p.144 and TI564 the fused version p.731/p.3912. **B**-Sequence alignment of derivates obtain after transduction. **C**-Growth curve of plasmid fused version vs transduce version.

